# HIF-1α promotes osteogenic-angiogenic coupling response of BMSCs cell sheets

**DOI:** 10.1101/2025.04.26.650808

**Authors:** Dan Zhang, Yonghui Teng, Yinli Huang, Wei Liu

## Abstract

**Background:** There is a critical need for management of vascularization in bone tissue engineering. The purpose of this study was to use HIF-1α-transduced BMSCs fabricated prevascularized osteogenic cell sheets and explore HIF-1α promoted osteogenic-angiogenic coupling response of BMSCs cell sheets in vitro.

**Methods:** HIF-1α was over-expressed by using a lentiviral vector, and transduced steadily in Wistar rats bone marrow mesenchymal stem cells(BMSCs). Real-time quantitative and western blot were performed to assess the expression level of HIF-1α. Then, HIF-1α/BMSCs were cultured to form osteogenic cell sheets(OCTs). ALP activity and Alizarin-red staining were performed respectively at day 14 and 21 to detect the characteristic of osteogenesis. Simultaneously, HIF-1α/BMSCs were induced to differentiate into endothelial-like cells(iECs) for 14 days, and flow cytometry was further detected the conversion rates of HIF-1α/BMSCs to iECs. Finally, iECs were seeded onto the OCTs to construct the prevascularized-osteogenic cell sheets (P-OCTs). In order to detect the role of HIF-1α involved in osteogenic-angiogenic coupling response of P-OCTs, Immunofluorescent staining for CD31 was performed on P-OCTs at 1, 3, 7, 14 days to check the formation of networks, and western blot of osteopontin(OPN) and osteocalcin (OCN) at 1, 7, 14 days to detect bone formation. Meanwhile, the none transduced BMSCs were as control.

**Conclusion:** BMSCs were transduced by Lenti-HIF-1α with optimal multiplicity of infection was 30(MOI=30), meanwhile, qPCR and western blot were performed to confirm this result. Next, the flow cytometric analysis results showed the conversion rate of BMSCs differentiated to iECs was 92.43%, which indicated BMSCs transduced by HIF-1α had a great superiority to differentiate into endothelial cells under experimental conditions. Then, ALP at day 14 and Alizarin-red staining at day 21 on OCTs showed an obvious osteogenic differentiation characteristic with more deep stained calcium nodules deposits than the control groups. Finally, we fabricated P-OCTs and observed iECs migrated reticulated fast and formed a large number of lumens and networks in experimental groups. At the same time, the results of Immunofluorescent staining for CD31 at day 1, 3, 7, 14 and osteogenic proteins expressions of OCN,OPN at day 1, 7, 14 showed that HIF-1α could promote osteogenic-angiogenic coupling response in P-OCTs significantly in vitro. All in all, the over-expressed HIF-1α of BMSCs cell sheets strategy can provide a new promising method for bone engineering, and we will detect further in vivo.

## Background

Large bone defects maily caused by trauma, tumor, inflammation are still a significant challenge in regeneration medicine due to the insufficient vascularization in engineered bone grafts^[1,2]^. Angiogenesis and osteogenesis are tightly coupled during bone development and regeneration. Insufficient vascularization induces lack of nutrients and oxygen which supplies a hypoxic-ischemic metabolic microenvironment for bone grafts causes low cell survival rate, slow growth of new bone, and bone grafts failure even more^[3,4,5]^. Recently, prevscularization cell sheet technology has been provided a promising strategy for bone engineering. Endothelial cells (ECs) are seeded on the 3D cell sheets with a certain density, then they migrate into cell sheets and form blood vessels gradually. Many studies^[6,7,8]^ have demonstrated that cell sheets-based prevascularizing system is beneficial for vascular network formation and enhance bone regeneration in vitro. However, the enhancement in vivo is still limited and sluggish resulting in precarious and insufficient prevascularized networks and poor connection with the host in the hypoxic-ischemic environment^[9,10,11]^. Therefore, there is a critical need for the management of vascularization in bone engineering at a hypoxic-ischemic metabolic microenvironment.

Hypoxia-inducible factor-1α(HIF-1α) is an active transcription factor that upregulates numerous genes at low oxygen conditions which involve in cell proliferation and differentiation, angiogenesis, glycolysis, and wound healing^[12]^. Angiogenic and osteogenic coupling is a complex and precise regulation process in bone healing, and a large number of studies show that HIF-1α plays an important role in this process. However, the expression of HIF-1α is restricted by the host while is overexpressed in cancers. With the development of biomedicine engineering, gene therapy has been investigated as a targeted therapy for bone healing. Sun et al.demonstrated HIF-1α-overexpressed exosomes could rescue the impaired angiogenic ability, migration, and proliferation by hypoxia-pretreated HUVECs in vitro and mediate cardioprotection by upregulating proangiogenic factors and enchancing neovessel formation^[13]^. Rachel Forster et al. found that gene therapy for peripheral arterial disease by increasing revascularisation was safe and efficacy^[14]^. Among genome engineering tools, Clustered Regularly Interspaced Short Palindromic Repeats(CRISPR) technology for ocular angiogenesis showed a huge potential by up-regulating the expression of HIF-1α^[15]^, et al. So, the usage of HIF-1α in biomedcine engineering is safe and efficient, which gives us some inspirations in bone engineering.

Therefore, we assumed that HIF-1α-overexpressed in BMSCs and used HIF-1α/BMSCs to construct osteogenic cell sheets(OCTs). Then, we induced HIF-1α/BMSCs differentiate into endothelial cells(iECs) and seeded them onto OCTs to fabricate prevascularized osteogenesis cell sheets(P-OCTs). Finally, we detected vascular networks formation and bone regeneration in vitro, in order to provide evidence for participation of HIF-1α in angiogenic and osteogenic coupling effects.

## Materials and Methods

### Ethical approval

The authors complied with ARRIVE guidelines, and all experiments adhered to the Principles for the Care of Laboratory Animals issued by the National Institutes of Health. Animal usage and experimental protocol were approved by the Experimental Animal Ethics Committee of Yinchuan Stomatology Hospital (YCKQLL2022018).

### rBMSCs harvest and culture

Three or four-week-old Wistar rats of 60-100g in weight were purchased from Jiangxi Zhonghongboyuan Biotechnology Company (Jiangxi province, China). The Wistar rats were anesthetized with diethyl ether and then killed by cervical dislocation. Rat bone marrow was aspirated from femoral medullary canal and supercentrifuged at 1500 r/min, then cultured in low-glucose Dulbecco’s modified Eagle’s medium(L-DMEM, Hyclone, USA) supplement with 10% fetal bovine serum(FBS, Hyclone, USA) and 2% antibiotics (100U/ml penicillin and streptomycin, Hyclone, USA), wihch was called L-DMED complete medium. BMSCs were cultured in a humidified incubator (Heraeus, Germany) of 5% CO_2_ at 37°C and the culture medium was replaced every 2 or 3 days. BMSCs were passaged with 0.25% trypsin/EDTA when adherent and confluent completely.

### HIF-1α transduction into BMSCs

The backbone vector pHBLV-CMV-MCS-EF1-ZsGreen-T2A-puro (Hanbio Biotechnology, Shanghai, China) was used for the reconstruction of a lentiviral vector containing HIF-1α. The Lenti-HIF-1α was made to overexpress using replication defective lentivirus that encoded green fluorescent protein(GFP), and BMSCs were transduced according to the product instruction. Then, a specific form of HIF-1α RNA was extracted according to the protocol provided in the HiScript II Q RT SuperMix for qPCR (+gDNA wiper)(R223-01,Vazyme) instructions. Real-time quantitative PCR was performed using the SuperStar Universal SYBR Master Mix(CW3360M, CWBIO) in the CFX Connect™ real-time system(Bio-rad, USA). The relative quantity of mRNA was calculated using the 2^-△△CT^ method. The sequences of primers designed by Generalbiol CO. Ltd.(Anhui, China) were listed in Table 1. The BMSCs group was as a control group, an empty lentivirus transduced BMSCs was employed as a negative control group(NC).

**Table 1.** Primers for RT-PCR analysis of HIF-1α.

### Western blot analysis

Cells were washed twice with cold PBS and lysed in 200μL RIPA cell lysis buffer(C1053, Applygen co.,Ltd., Beijing, China) for 10-20 minutes, and then scraped into microfuge tubes and centrifuged for 10 minutes at 12000 r/min. Collected the supernatant and the protein concentration was measured by BCA Protein Assay Kit (E-BC-K318-M, Elabscience,Wuhan, China). Subsequently, equal amounts of cell lysates were separated by SDS-PAGE gel(A1010,Solarbio) and transferred to polyvinylidene fluoride membranes(PVDF,IPVH00010,Millipore). The membranes were blocked with 3% skim milk(P1622, Applygen Co.,Ltd., Beijing,China) in Tween Tris-buffered saline(TTBS)(0.5%, Tween-20) for 1 hour. The blocked membranes were incubated with primary antibodies (Rabbit Anti HIF-1α,bs-0737R,Bioss,1/1000) overnight. After washing in TTBS, membranes were then incubated with secondary antibodies(HRP conjugated Goat Anti-Rabbit IgG (H+L), GB23303, Servicebio,1/2000) for 2 hours at room temperature. Images for western blot analysis were visualizes by automatic chemiluminescence image analysis system (Tanon-5200,Shanghai Tanon Life Science Co.,Ltd, China). Relative protein levels were calculated as ratio level of protein of interest to β-actin in each sample.

### Differentiation of HIF-1α/BMSCs into endothelial-like cells

The second generation of HIF-1α/BMSCs was seeded into six-well plates at a density of 3×10^4^ cell/cm^2^ and cultured with L-DMED complete medium in a humidified atmosphere of 5% CO_2_ at 37°C. When HIF-1α/BMSCs reached confluence, the culture medium was replaced by M199 medium(Hyclone, USA) plus 10% FBS, vascular endothelial growth factor(VEGF, 10μg/L) and basic fibroblast growth factor (bFGF, 2μg/L), which referred as M199 complete medium. HIF-1α/BMSCs were differentiated for 14 days, while HIF-1α/BMSCs cultured without VEGF and bFGF were as a control.

### Flow cytometric analysis of endothelial-like cells

Flow cytometric analysis was used to detect whether the i-ECs had the endothelia specific phenotype PECAM-1(CD31). After differentiated culture for 14 days, HIF-1α/BMSCs were trypsinized (0.25% trypsin/EDTA), and added into an Eppendorf tube with a density of 1×10^6^/mL, then incubated with the monoclonal antibody CD31(25-0310-82, Thermo Scientific, China) for 30 min at 4°C. While non-differentiated HIF-1α/ BMSCs group served as a control. Afterwards, the samples were performed on the FACSVerse flow cytometer (BD,USA)and analyzed the CD31-positive expression.

### Fabrication of osteogenesis cell sheets

HIF-1α/BMSCs were seeded into the six-well plate with a density of 1×10^5^/cm^2^ and cultured in osteogenesis medium which contained 12% FBS, 50mg/L ascorbic acid, 10nM dexamethasone, and 10 mM β-glycerophosphate for 21days to fabricate osteogenesis cell sheets(OCTs). Alkaline phosphatase (ALP) at day 14 and Alizarin red-S staining(ARS) at day 21 were performed to characterize the osteogenic properties of OCTs. At the same time, NC group formed OCTs served as a control.

### Fabrication of prevascularized osteogenesis cell sheet

To fabricate prevascularized osteogenesis cell sheets(P-OCTs), i-ECs were seeded onto the osteogenesis cell sheets surface with a density of 5×10^4^/cm^2^ and cultured with a mixed medium which concluded M199 complete medium and osteogenesis medium(1:1,v/v). The culture medium was replaced every 3 days. The NC group formed P-OCTs served as a control.

### Immunofluorescent staining of prevascularized osteogenesis cell sheet

The fluorescent microscope was used to detect the i-ECs migration in the OCTs. After cultured for 1,3,7,14 days, the P-OCTs was washed with phosphate-buffered saline(PBS), fixed in 4% paraformaldehyde for 15 minutes, and then blocked in 5% goat serum-PBS buffer solution for 1 hour at room temperature. A primary antibody rabbit anti CD31 (GB11063-2, Servicebio, dilution 1/200) in 1% bovine serum albumin(BSA)-PBS was added to the samples and incubated overnight at 4%. After washing with PBS, a secondary antibody Cy3 Goat Anti-Rabbit IgG(H+L) (AS007, ABclonal, dilution 1/200) in 1% BSA-PBS buffer was added and incubated in the dark for 1 hour at room temperature. Finally, the cell nuclei were counterstained with DAPI((KGA215-50, keygentec. Jiangsu, China) for 5 min and rinsed three times with PBS in the dark. The fluorescent staining images were captured by confocal microscopy (Research invested system microscope, IX71, Olympus).

### Osteogenesis proteins expression of prevascularized osteogenesis cell sheet

Osteogenesis proteins were extracted and electroblotted as above-mentioned after P-OCT cultured for 1,7,14 days. The same procedure was followed on reaction products of the following antibodies, including to osteocalcin(OCN)(Rabbit Anti OCN, A20800, Abclonal, 1/1000), osteopontin(OPN)(Rabbit Anti OPN, YT3467, Immunoway, 1/1000).

### Statistical analysis

All data were express as mean ± standard deviation. The variance(ANOVA) and Tukey post hoc tests were performed to analyze the results by using SPSS17.0 software, where a significant difference was considered if the *P* value was less than 0.05.

## Results

### HIF-1α transduction

Through a set of preliminary experiments using various doses of lentivirus, we found the optimal multiplicity of infection(MOI=30). After transduction, BMSCs fluoresced green under inverted fluorescence microscopy, showing efficiency of transduction of approximately 90%(Figure1A, B). HIF-1α mRNA and protein expression were regulated in the target groups compared with the control group and the negative control group(virus without HIF-1α, NC) by qPCR and western blot(Figure 1C, D,E). The quantitative results showed a significant HIF-1α mRNA and HIF-1α protein expression in HIF-1α transduced group between the control group and the negative control group(*: the control group vs HIF-1α transduced group; #: NC vs HIF-1α transduced group), while the control group and NC had no statistical significance. (*P*<0.05).

**Figure 1.**
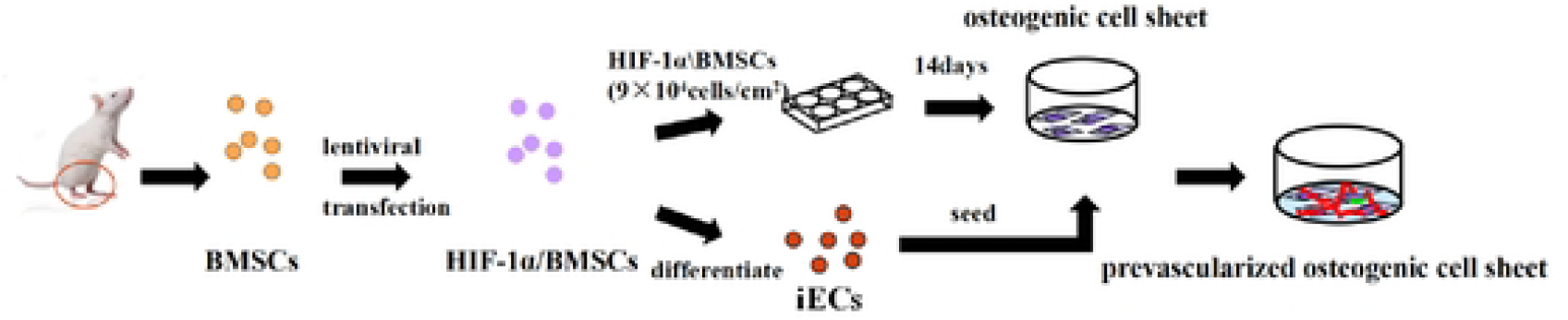
The schematic shows the whole procedure of the experiments. First, the rat bone marrow mesenchymal stem cells(BMSCs) were harvested and cultured. Then, the second generation of BMSCs were transduced by Lenti-HIF-1α. HIF-1α/BMSCs were cultured in osteogenesis medium to fabricate osteogenesis cell sheets(OCTs). On the other hand, HIF-1α/BMSCs were induced to differentiate into endothelial-like cells(iECs). Finally, iECs were seeded onto osteogenesis cell sheets to construct the prevascularized osteogenesis cell sheets(P-OCTs).

### iECs morphology and characterization of iECs

Figure 3 showed cell morphology and flow cytometry results of iECs. It seemed that iECs were more oval, short and swirling grew, while HIF-1α/BMSCs were spindle-shaped spreading and long(Figure2A). Flow cytometry results showed that HIF-1α/BMSCs without induced had only 0.81% CD31 positive expression, but after differentiated into iECs by endothelial culture medium which expressed 92.43%(Figure3B,C). Quantitative results(Figure3D) showed a significant difference in differentiated group(iECs group) and non-differentiated group(HIF-1α/BMSCs group) by counting CD31 positive expression (*p*<0.05.), which indicated BMSCs transduced by HIF-1α had a great superiority to differentiated into endothelial cells under experimental condition.

**Figure 2.**
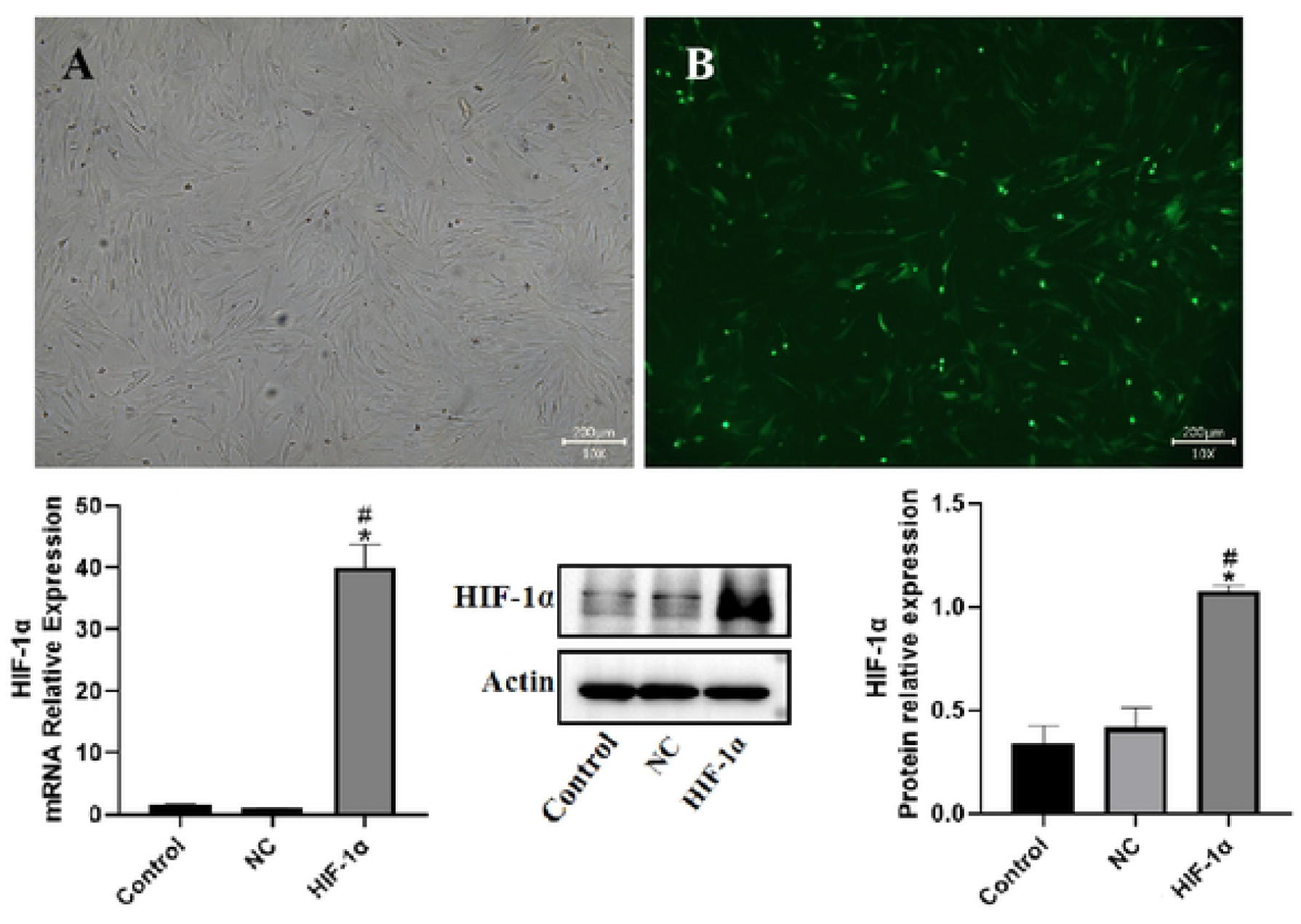
The microscopic and fluorescence microscopic morphology of HIF-1α/BMSCs (A, B), which performed a well transfection. HIF-1α mRNA expression(C),Western Blot(D), and HIF-1α protein expression(E) between control group, negative control group(virus without HIF-1α, NC) and HIF-1α induced group. Quantitative results showed the tremendous expression of HIF-1α in HIF-1α induced group among those two groups (p<0.05). Scale bars=200μm (A,B).

**Figure 3.**
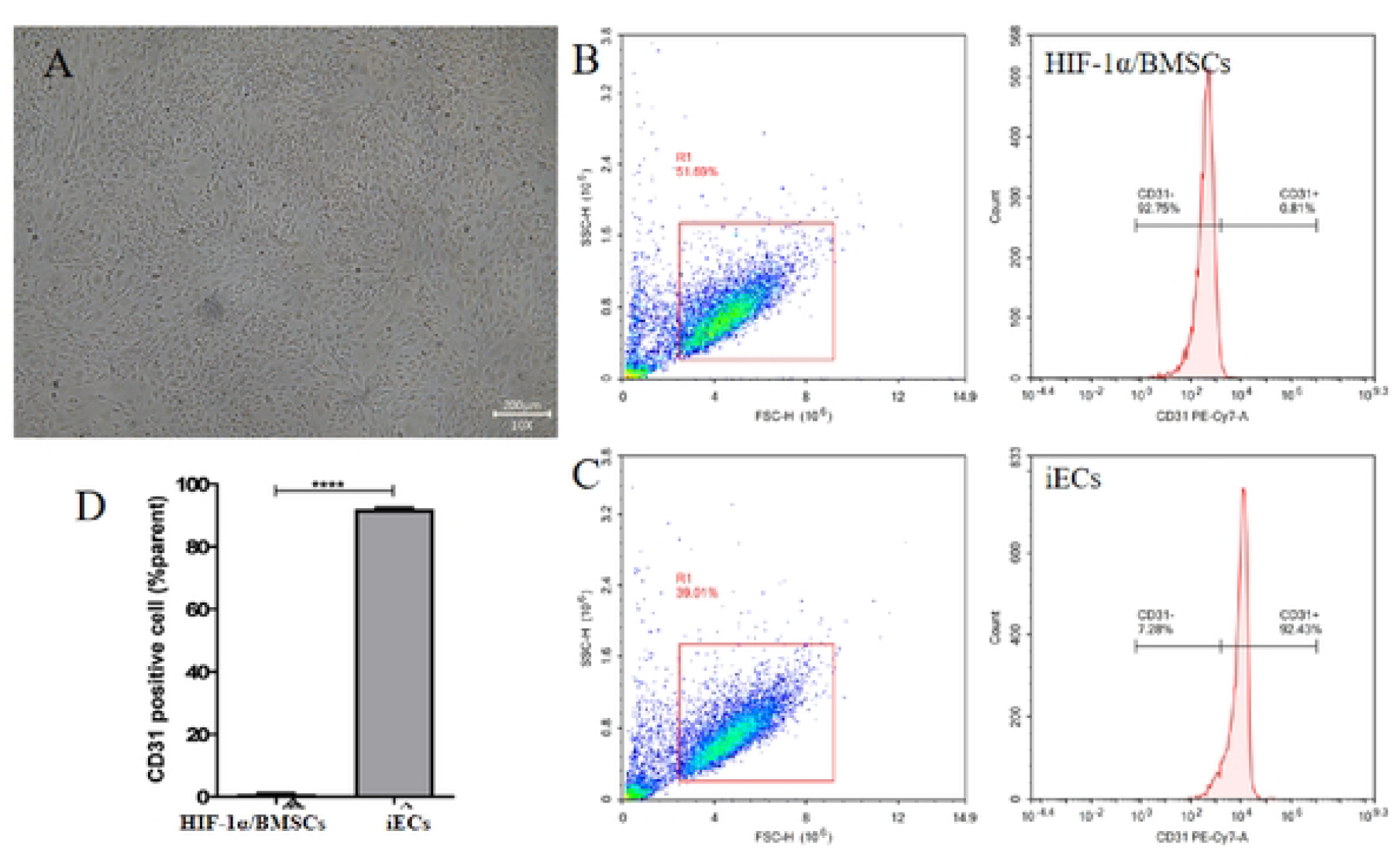
Cell morphology of induced endothelial cells(iECs)**(A)**.iECs were oval and short, while HIF-1α/BMSCs were spindle-shaped spreading and long(Fig.2). Flow cytometric analysis of CD31 of HIF-1α/BMSCs group(nondifferentiated group) **(B)** and iECs group(differentiated group) **(C)**.Quantitative results**(D)** showed a significant difference between two groups which indicated BMSCs had the ability to differentiated into endothelial cells under experimental condition. Scale bars=200μm.

### Characteristics of osteogenesis cell sheets

HIF-1α/BMSCs were cultured in osteogenesis medium for 21 days to fabricate osteogenesis cell sheets(OCTs) (Figure4A) which performed a dense and turbid film with the edges curled inward with time increased over 21 days(Figure4B), and sometimes it could be lifted up by point forceps. ALP(Figure4C) at day 14 and Alizarin red-S staining(Figure4D) at day 21 showed both two kinds of OCTs had osteogenic differentiation under the same condition, but HIF-1α/BMSCs formed OCTs had an obvious and tremendous osteogenic differentiation characteristic with massive stained calcium nodules deposited than the control group. Meanwhile, the quantitative results(Figure4E, F) showed a significant osteogenic characteristic deviation between HIF-1α/BMSCs formed OCTs and the negative control group(NC), which indicated HIF-1α could promote the potential for osteogenic differentiation.

**Figure 4.**
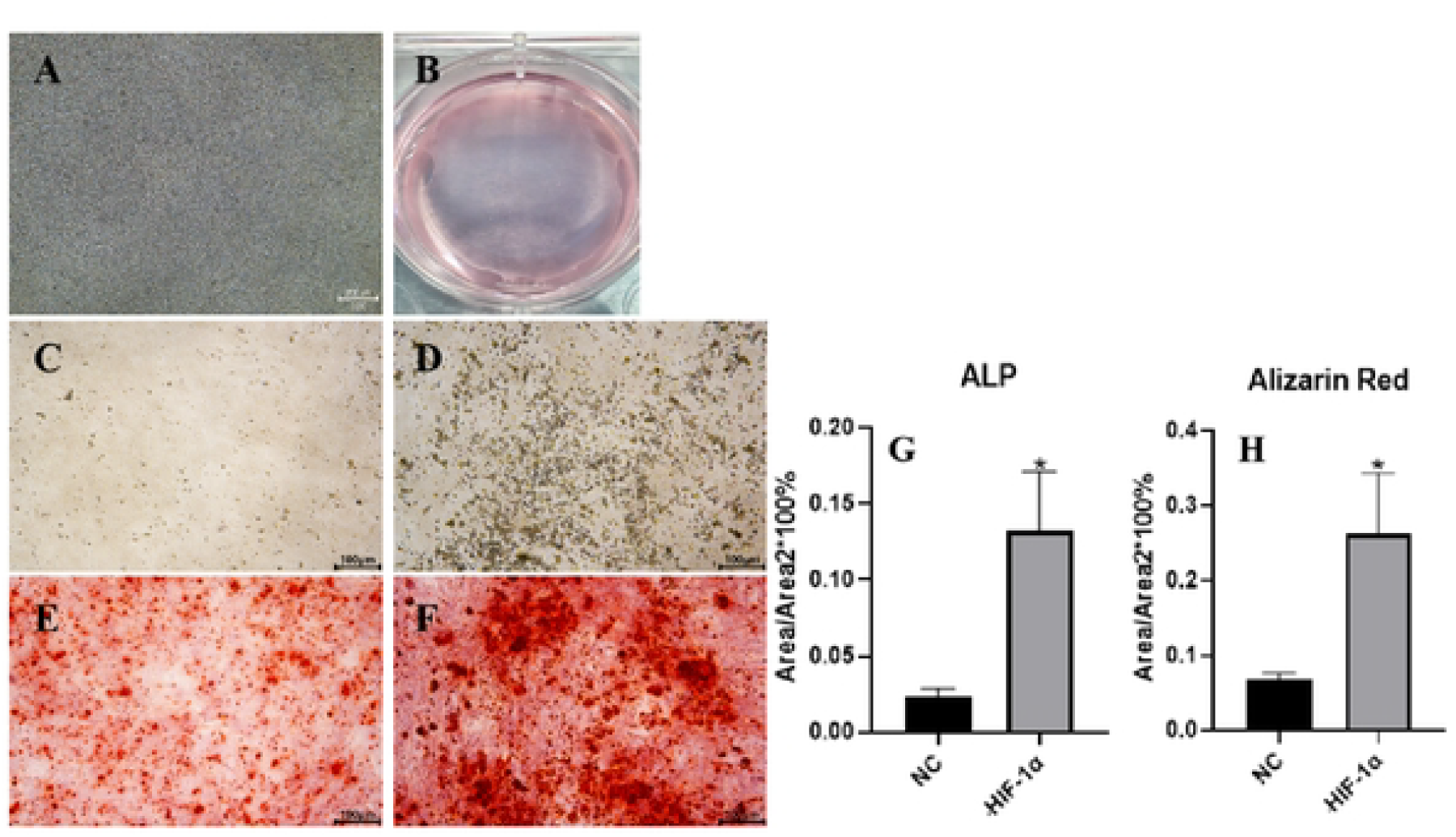
The microscopic morphology and view of osteogenesis cell sheets(OCTs)(A,B), which performed a dense and turbid film with the edges curled inward. ALP at day 14 and Alizarin red-S staining at day 21 showed osteogeneic differentiation characteristic of the OCTs. Quantitative results(G,H) showed a significant osteogenic differentiation differences between two groups (*p*<0.05). C,E: NC group; D,F: the experimental group. Scale bars=200μm(A) and 100μm (C,D,E,F).

### Immunofluorescent staining for CD31 of pre-osteogenesis cell sheets

When iECs seeded onto OCTs to construct P-OCTs, the microscopic morphologies became so different than before(Figure5). With time increased, P-OCTs showed grid multilayers and became more and more compact. In order to detect the role of HIF-1α involved in osteogenic-angiogenic coupling response of P-OCTs, Immunofluorescent staining for CD31 was performed to detect the networks formation of iECs on OCTs. As shown in Figure6, at day 1, iECs adhered and proliferated massively on OCTs, then they migrated fused into OCTs which showed a random arrangement and continued to form a large number of alignment with each other. The experimental group showed a large number of iECs proliferation than the control group(NC), and with a more and quick cell-to-cell alignment. it seemed that the vacuoles structures were formed in experimental group(Figure6A). At day 3, with the multiplication of iECs, they migrated reticulated and formed a large number of lumens in experimental group (Figure6B), while the NC group began to formed the vacuoles structures. At day 7, the lumens began to connect and formed stable tubulose vascular networks in experimental group (Figure6C). on the other hand, NC group began to form the lumens. At day 14, the lumens became thicker and compact(Figure6D). The experimental group formed a more stable and a large quantity of networks in the OCTs, while formed some sparse lumens in the NC group. Quantitative reselts showed a significant difference in vascular networks formation among experimental group and NC group(p<0.05) by counting CD31 positive expressing lumens at day 1,3,7,14. The networsk formation was increasing by time which was highest at day 7, and it seemed that the experimental group was more than twice as NC group. With time increased, the lumens had a sharp decrease in NC group, but still kept a relative stability in experimental group. At last time, the quantities of lumens in experimental group were more than three times as NC group.

**Figure 5.**
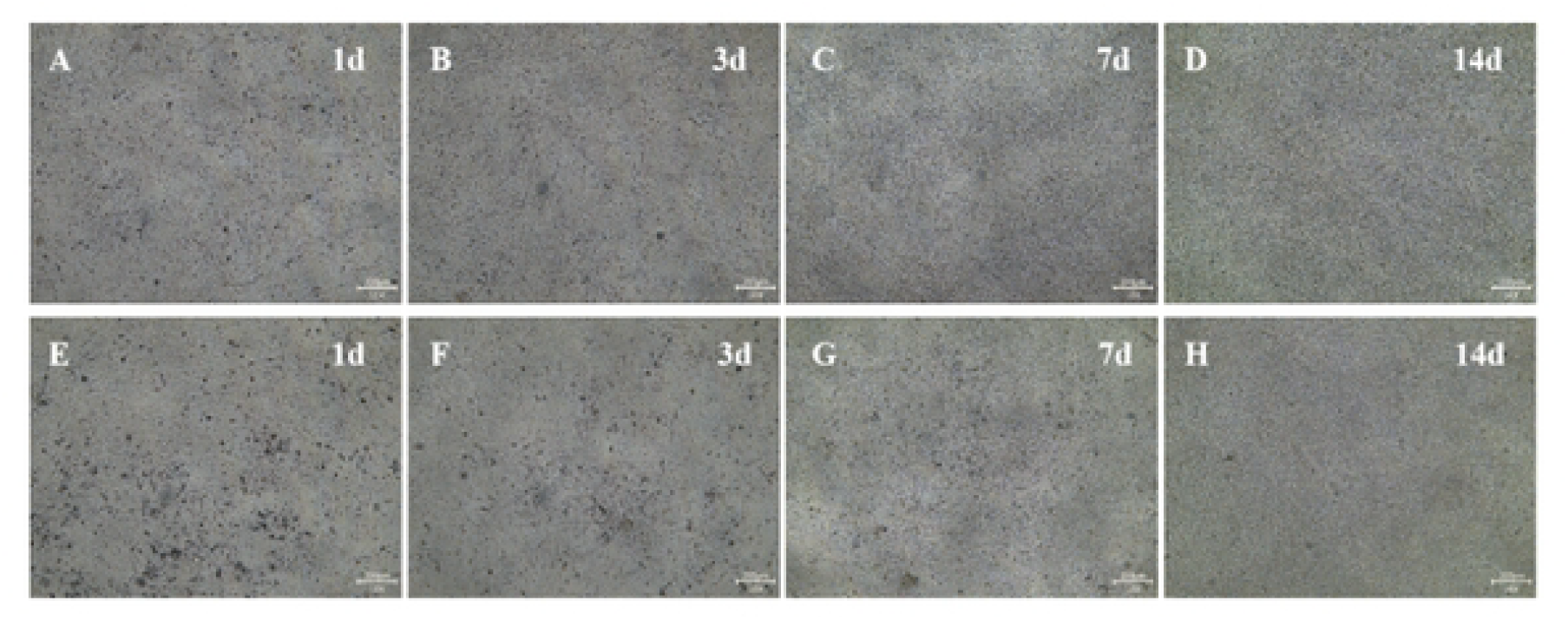
The microscopic morphology of prevascularized osteogenesis cell sheets(P-OCT). The control group(NC): A,B,C,D; the experimental group:E,F,G,H; at day 1, 3, 7, 14. Scale bars = 200μm.

**Figure 6.**
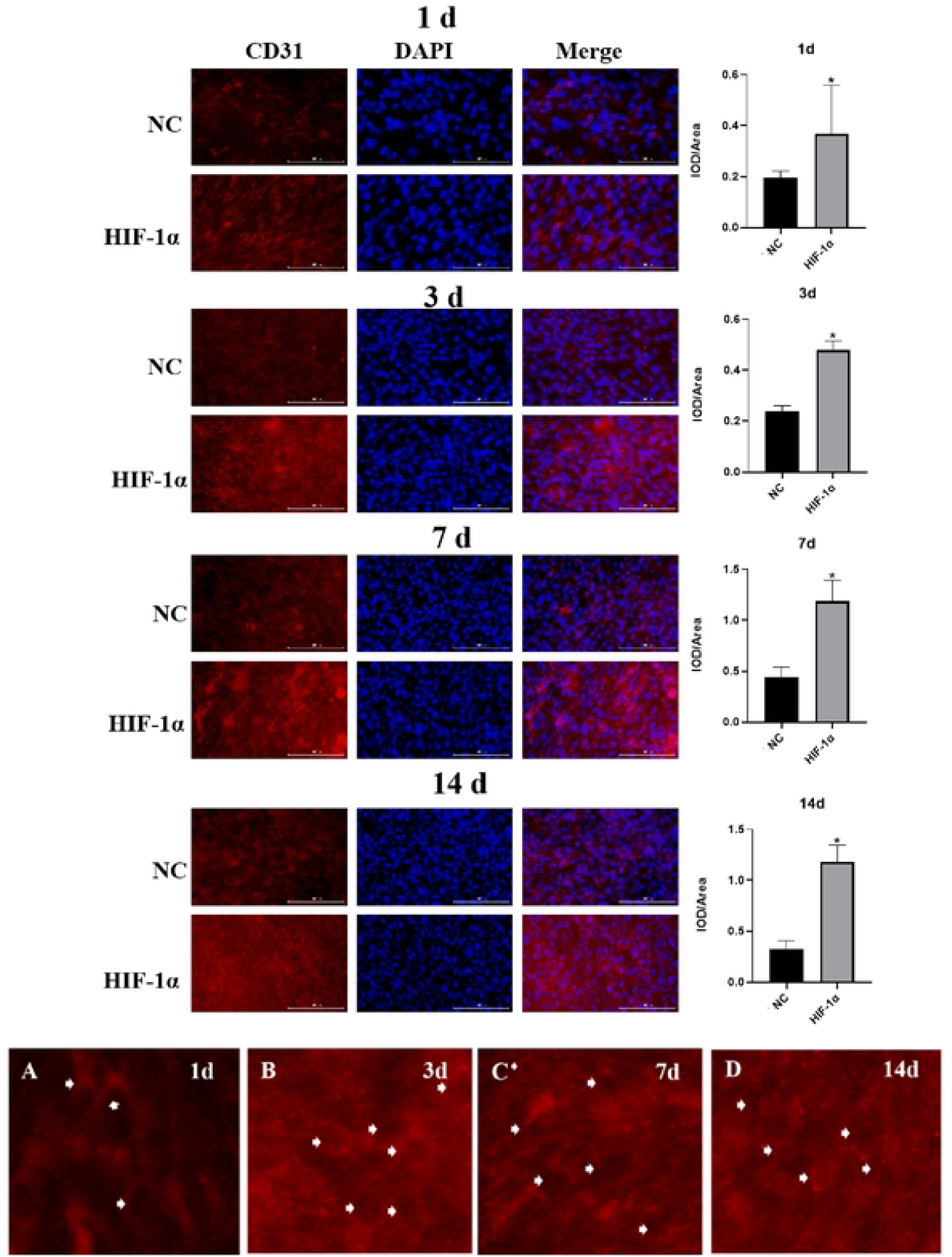
Immunofluorescent images of CD31 for prevascularized osteogenesis cell sheets and quantitative results at day 1, 3, 7, 14 (p<0.05). The networks formation in experimental group was increasing by time which was highest at day 7, and still kept a relative stability at day 14, but the lumens had a sharp decrease in NC group at day 7. A, B, C, D: the magnified picture of experimental group at day 1, 3, 7, 14. (Scale bars=100μm, Red: CD31 antibody, Blue: DAPI.)

### Analysis of osteogenesis proteins expression

HIF-1α has profound effects on osteogenesis in OCTs. To further explore the role of HIF-1α involved in osteogenic-angiogenic coupling response of P-OCTs, we analyzed osteogenic proteins expressions of OCN,OPN by western blot(Figure 7). At day 1, the expression of OCN in NC group was highest. With time increased, OCN had a down-regulated expression and stayed at a level at day 14. Meanwhile OCN in experimental group was up-regulated and highest at day 7, then down-regulated slowly at day 14. All in all, the expressions of OCN in experimental group were higher than NC group at any time point(*P*<0.05). On the other hand, the expressions of OPN were highest at day 1, and had a quick down-regulation at day 7. With time increased, the expression leveled off at day 14. On the whole, the expressions of OPN in the experimental group were higher than NC group at any time point(*P*<0.05).

**Figure 7.**
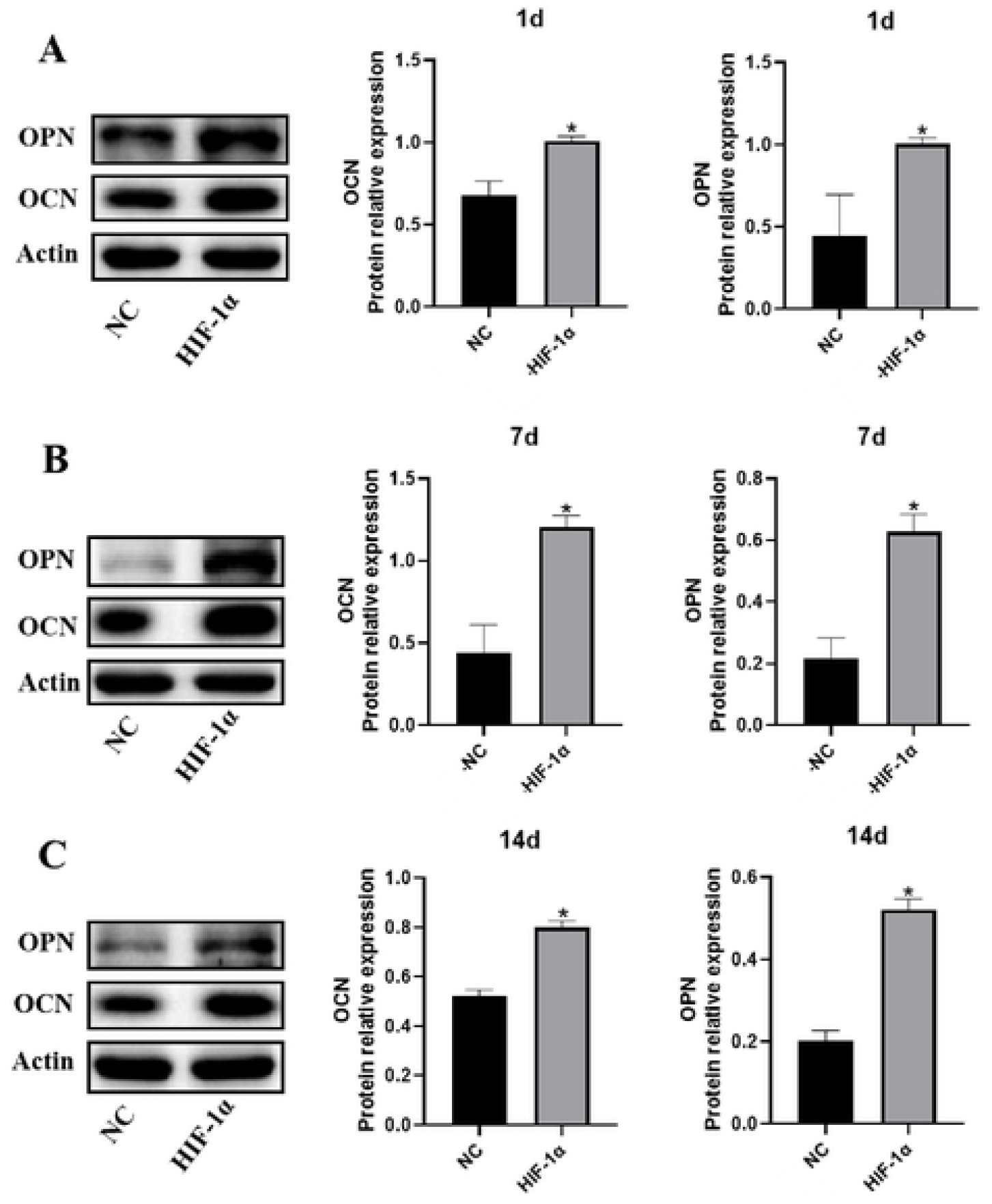
Western Blot of OCN and OPN at day 1, 7, 14. In experimental group OCN expression was highest at day 7 and OPN at day 1; in NC group, OCN and OPN expressions were both highest at day 1, but lower than experimental group at any time point(*P*<0.05).

**Figure 8.** Full-length blots are presented in Supplementary.

## Discussion

Sufficient and stable vascularization plays an important role in tissue-engineered bone constructs for optimal cell survival and implants integration, but which is limited and complicated in hypoxic-ischemic conditions^[16]^. At this critical point, HIF-1α is an active transcription factor which plays an important role in angiogenesis and osteogenesis by up-regulating numerous genes and relevant factors^[17]^. Therefore, we assumed that HIF-1α-overexpressed in BMSCs to construct prevascularized osteogenic cell sheets in vitro, and we detected the effect of HIF-1α in angiogenic and osteogenic coupling response.

For the aspect of angiogenesis, first of all, we used endothelial culture medium contained VEGF and bFGF to induce HIF-1α/BMSCs for 14 days, then the flow cytometric analysis showed over 92.43% CD31 positive expressions. However, in our previous study^[18]^, the conversion rate was just 35.1%, which indicated BMSCs transduced by HIF-1α had a great superiority to differentiate into endothelial cells under experimental conditions. Then we seeded iECs onto osteogenesis cell sheets(OCTs) to fabricate prevascularization. We observed iECs migrated reticulated fast and formed a large number of lumens and networks in experimental groups. On the other hand, we examined ALP at day 14 and Alizarin red-S staining at day 21 of OCTs to detect whether HIF-1α could improve the ability of osteogenesis in vitro. The results showed an obvious osteogenic differentiation characteristic with more deep stained calcium nodules than control groups(*p*<0.05). Furthermore, we checked the expressions of OCN, OPN by western blot, which showed statistical significance with control groups at any time points(*P*<0.05). All in all, we concluded that HIF-1α-overexpressed in BMSCs could promote osteogenic-angiogenic coupling response by constructing prevascularized osteogenic cell sheets in vitro.

HIF signaling pathways induced by hypoxia were involved in angiogenesis and osteogenesis by modulating vital genes, such as VEGF and EPO, and was a crucial regulator that influenced final fate of bone regeneration^[19,20]^. Studies have been demonstrated that knockdown HIF-1α in vivo or in vitro significantly impaired bone generation and osteogenesis of periosteum-derived stem cells, indicating the indispensability of HIF-1α in bone regeneration under hypoxia^[21]^. In our study, we adopted 10μg/L VEGF and 2μg/L bFGF in M199 media to culture HIF-1α/BMSCs and construct prevascularization which showed a positive function under HIF-1α’s regulation. It has been widely reported that HIF-1α/VEGF signaling pathway plays a key role in cell proliferation, migration, angiogenesis, and osteogenesis in anoxic environment^[22-24]^. And recent studies have investigated the coupling of HIF-1α-driven bone formation with angiogenesis induced by VEGF upregulation in vivo^[25,26]^, and studies have reported that activation of VEGF/AKT/mTOR signaling pathway could promote angiogenesis and differentiation of BMSCs ^[22]^. Meanwhile, bFGF played a major role in tissue repair and regeneration with a recognized role in epithelial and mesenchymal cell proliferation as well as a putative function in angiogenesis^[27]^. Furthermore, studies showed that FGF signaling could inhibit apoptosis and promote angiogenesis via HIF1α-meidated mechanism^[28]^. On the other hand, we fabricated OCTs by confluent culturing HIF-1α/BMSCs for 21days. As we all known, cell sheet technology takes advantages of its densely populated microenvironment which determines cell fate and function^[29]^. OCTs with cell sheets’ superiority that retain deposited extracellular matrix(ECM), cell adhesive proteins and cell-cell and cell-matrix interactions generated during confluent culture^[30,31]^. Studies indicated the retention of active components in ECM, such as TGF-β1, BMP-2, collagen type I, tenascin, Emilin, which had high osteoconductive capacity^[32,33]^. In our study, we found when iECs seeded onto OCTs formed by HIF-1α/BMSCs, they migrated, rearranged and formed vascular networks on OCTs efficiently. The microenvironment and crosstalk between endothelial cells and bone-forming cells are essential for successful bone regeneration, due to the endothelial cell matrix facilitates osseointegration and mineralization, in turm, osteogenic cells secrete paracrine factors and stimulate angiogenesis^[34,35]^. Studies found the activation of TGF-β and Notch signaling pathway in the endothelial cells during both contact and indirect co-cultivation with osteoblasts, which promoted bone formation^[36]^.

## Conclusion

HIF-1α over-expressed in BMSCs shows a great potential in angiogenic and osteogenic differentiation. On the one hand, HIF-1α/BMSCs had a great superiority to differentiated into endothelial cells under experimental condition, which could provide a steady flow of endothelial precursor cells for constructing prevascularization. In addition, iECs seeded onto osteogenic cell sheets showed a quick and extensive connection to each other and formed more stable tubulose vascular networks. On the other hand, HIF-1α/BMSCs showed a significant osteogenic differentiation characteristic, which had more deep stained calcium nodules deposits than the control groups. All in all, HIF-1α plays an important role in angiogenic and osteogenic coupling response of BMSCs sheets in vitro, which provides a promising strategy for bone engineering.

## Acknowledgments

The authors declare that that they have not use AI-generated work in this manuscript.

## Availability of data and meterials

Not applicable.

## Authors’ contributions

DZ, YT, YH, and WLiu participated in the acquisition of data, analysis and interpretation of data, and writing of the manuscript. DZ participated in the study design, analysis and interpretation of data, and writing of the manuscript. All authors read and approved the final manuscript for publication.

## Ethics approval and consent to participate

The project named the Effect and Mechanism of HIF-1α on osteogenic and angiogenic coupling reaction of prevascularized BMSCs cell sheets was approved by the Nature Science Foundation of Ningxia Province (No.2023AAC03880). The Animal Ethics Committee of Yinchuan Stomatology Hospital approved all the experimental animal procedures and sample collections at September 9, 2022 and the approval nomber was YCKQLL2022018.

## Table caption

